# The Lipid-binding PX Domain of RRC-1 (ARHGAP32/33) is Required for Optimal Assembly and Function of Integrin Adhesion Complexes at the Muscle Cell Boundary in C. elegans

**DOI:** 10.64898/2025.12.12.694024

**Authors:** Sara Sagadiev, Isabel Martin, Nahum Arefeayne, Richard (Yuchu) Wang, Robert Hudson, Olga Mayans, Hiroshi Qadota, Guy M. Benian

## Abstract

Integrin adhesion complexes (IACs) are a network of many proteins that serve as anchors of the cell to the extracellular matrix (ECM). In muscle, IACs located at costameres, also serve to transmit the force of muscle contraction to the outside of the cell. We have reported that IACs, which are found at the bases of dense bodies and M-lines, and at muscle cell boundaries (MCB) in *C. elegans* muscle, require the RacGEF PIX-1 for their proper assembly or maintenance. We have reported that a RacGAP for the PIX pathway is RRC-1, is in a complex with PIX-1, and that RRC-1 is required for assembly or maintenance of IACs at MCBs. Our previous studies suggested that RRC-1 might be associated with the muscle cell membrane, and here we present evidence that this occurs via its PX domain, a domain that is known to bind to membrane phosphoinositides (PIPs). We predict the existence of a PX domain based on bioinformatic analysis and AlphaFold3, which includes conserved residues characteristic of most PX domains and a PIP binding site. This region of RRC-1 binds to phosphoinositides in vitro. Analysis of a nematode strain that has an in-frame deletion of the PX domain, indicates that normal localization of RRC-1 to the MCB requires both its PX domain and the PIX scaffold protein GIT-1. Lastly, we show that the overexpression of the full length RRC-1, but not RRC-1 with an in-frame deletion of its PX domain, results in reduced accumulation of IAC components and reduced whole animal movement. Our study highlights the importance of RRC-1’s lipid interactions at the cell membrane for proper assembly and function of IACs in *C. elegans* muscle.

**Article Summary:** Integrin adhesion complexes (IACs) facilitate the transmission of force of muscle contraction to the outside of the cell. IACs, found at the bases of dense bodies and M-lines, and at muscle cell boundaries (MCB) in *C. elegans* muscle, require the PIX-1 (a RacGEF) signaling pathway for their assembly or maintenance. A RacGAP for the PIX pathway is RRC-1. Here, we show that RRC-1 has a PX domain that binds to membrane phosphoinositides. We also show that normal localization of RRC-1 to the MCB requires both its PX domain and the PIX scaffolding protein GIT-1.

## Introduction

Integrin adhesion complexes (IACs), often referred to as focal adhesions, consist of the transmembrane α/β heterodimeric integrin and hundreds of other proteins, intracellular proteins associated with a cytoplasmic tail and extracellular matrix (ECM) proteins associated with the extracellular portion of integrin (Anthis and Campbell 2011, Alexia, Zareno et al. 2014). Dynamic assembly and disassembly of IACs are crucial for migrating cells during development, inflammation and cancer, and for stationary cells to attach to the ECM (Horton, Byron et al. 2015). In striated muscle, which includes both skeletal and cardiac muscle, myofibrils at the periphery of the cell are attached to the cell membrane and ECM via “costameres”, muscle-specific IACs (Ervasti 2003). Costameres function to anchor the muscle cell to the ECM and transmit the force of muscle contraction to the outside of the cell. In the genetic model organism *C. elegans*, the major striated muscle is localized to the body wall and through it cyclical contraction and relaxation permits the whole nematode to move (Benian and Epstein 2011). The organization of its striated muscle is like other animals, where the thin filaments attach to Z-disk-like structures, called dense bodies, and thick filaments are cross-linked at M-lines. In *C. elegans* muscle, specific IACs are located at M-lines, dense bodies, and muscle cell boundaries (MCB) with the base consisting of integrins and a core set of proteins along with site-specific proteins(Gieseler, Qadota et al. 2017, Qadota, Matsunaga et al. 2017). Our laboratory was the first to report that the PIX signaling pathway has an important role in striated muscle (Moody et al. 2020). The *C. elegans* protein PIX-1 is required for the assembly or stability of IACs at the MCB. PIX is a RacGEF that activates the small GTPases Rac and Cdc42 by promoting the exchange of GTP for GDP (Mosaddeghzadeh and Ahmadian 2021). GTP-Rac binds to and activates effector protein kinases (called PAKs), which then phosphorylate unidentified substrates. In some still unknown way, activation of this pathway promotes the assembly of IACs. Loss of function mutations in any component of the known PIX pathway results in a similar phenotype—loss of assembly of IACs at the MCB. Moreover, either deficiency or overexpression of PIX-1 results in disrupted MCBs and decreased muscle function, indicating that the level of PIX-1 needs to be tightly controlled.

Rho family GTPases (Rho, Rac, Cdc42) cycle between active (GTP bound) and inactive (GDP bound) states. In the PIX pathway, activation occurs by PIX. Inactivation occurs via so called GTPase-activating proteins (GAPs), which promote the hydrolysis of GTP to GDP. We have reported identification of the first RacGAP for the PIX pathway in any organism or cell type, called RRC-1 (Moody et al., 2024). Loss of function of *rrc-1* results in reduced localization of IACs to the MCB, disorganization of M-lines and dense bodies, and reduced whole animal locomotion. Multiple lines of genetic and cell biologic evidence suggest that RRC-1 is part of the PIX-1 pathway: (1) each single mutant results in a similar defect at the MCB; (2) PIX-1 and RRC-1 each localize to the MCB; (3) localization of RRC-1 to the MCB depends on PIX-1 and vice versa; and (4) RRC-1 exists in a complex with PIX-1.

Several pieces of data suggest that RRC-1 might be membrane-associated (Moody et al., 2024). First, in addition to localizing to the MCB, RRC-1 is also localized at what seems to be the bases of the M-lines and dense bodies but in a fuzzy and indistinct manner reminiscent of the localization of the ECM protein UNC-52 (perlecan). Second, to demonstrate coIP of RRC-1 with PIX-1, it was necessary to use a small amount of the ionic detergent, deoxycholate, in addition to a non-ionic detergent to extract RRC-1 from worms. Third, the human orthologs for RRC-1, ARHGAP32 and ARHGAP33, have predicted phox homology (PX) domains. PX domains are known to bind to phosphoinositides (PIPs) and target proteins to membranes (Chandra and Collins, 2019). However, in our previous report on RRC-1 (Moody et al., 2024), we could not predict a PX domain in RRC-1 using PFAM and ProSite programs. Here, we report that we can now predict a PX domain by sequence, and by homology modeling, and that indeed this PIX domain binds to PIPs. By generating an in-frame deletion of the PX domain, we find that it is required for normal worm locomotion. Moreover, optimal localization of RRC-1 to the MCB requires both its PX domain, and the PIX-1 scaffold protein GIT-1. Finally, we demonstrate that overexpression of full-length but not PX-deleted RRC-1 interferes with the localization of IAC components to the MCB and normal whole animal locomotion.

## Methods

### Prediction of the existence of a PX domain in RRC-1 and generation and analysis of its 3D-model

The 3D-fold of the PX domain from RRC-1 (residues 41-146; UniProtKB Q20498-1) was modelled using AlphaFold3 (Abramson et al. 2024). The predicted model was compared to all protein entries at the RCSB Protein Data Bank (https://www.rcsb.org/) using the DALI server (http://ekhidna2.biocenter.helsinki.fi/dali/; Holm, 2022). The top ten experimental structures identified in this way were structurally aligned to the PX-RRC-1 domain using USFC Chimera version 1.19 (https://www.cgl.ucsf.edu/chimera/; Pettersen et al. 2004). Matchmaker function in default settings (chain pairing: bb, Needleman-Wunsch using the BLOSUM-62, ss fraction: 0.3, gap open (HH/SS/other) 18/18/6, extend 1, ss matrix: (O, S): −6 (H, O): −6 (H, H): 6 (S, S): 6 (H, S): −9 (O, O): 4, iteration cutoff: 2) (Meng et al. 2006). A structure-based multiple sequence alignment was then generated from the superimposed models using USFC Chimera’s Match->Align function in default settings and alignment conservation scores were calculated using the built-in conservation annotation function, based on sequence identity and similarity of physico-chemical properties (Pettersen et al. 2004; Meng et al. 2006). The percentage of sequence identity in the Match-> Align function was calculated with the reference sequence set to the RRC-1 PX domain.

#### Expression and purification of GST, and GST-RRC-1 (1-150)

To create a plasmid for bacterial expression of GST-RRC-1(1-150aa), the coding sequence for residues 1-150 was amplified by PCR using as template a full-length *rrc-1* cDNA (Moody et al., 2024), and the following primers:

Rrc-1 anti-N1:BamHI-ATG-
GCGGGATCCATGGAAGGCATCGAGGAATCA
Rrc-1 anti-N2:-150aa-XhoI
CGCCTCGAGTTAGCCACGAGAGTCAATTTCTAA

The amplified fragment was cloned into the BamHI and XhoI sites of pGEX-KK-1 and a clone was identified to be error-free by Sanger sequencing. To create a plasmid for bacterial expression of 6xHis-RRC-1(36-150aa), the coding sequence for residues 36-150 was amplified by PCR using as template a full-length *rrc-1* cDNA (Moody et al., 2024), and the following primers:

His-RRC-1-PX-1: NcoI-36aa-
GCGCCATGGCGTTCCATTATTCGTCTGTTGAATTG
His-RRC-1-PX-2: −150aa-STOP-XhoI
CGCCTCGAGTTAGCCACGAGAGTCAATTTCTAAAAATTTC
The amplified fragment was cloned into the NcoI and XhoI sites of pETM11 and a clone was identified to be error-free by Sanger sequencing.

pGEX-KK1, and pGEX-KK1-RRC-1 (1-150) were transformed into E. coli Rosetta 2 (DE3) and used to produce proteins using procedures described in Matsunaga et al. (2024). A Bradford Assay (BioRad, Inc.) was used to determine the protein concentrations, and the proteins were stored on ice until used.

### Lipid protein overlay assay with a PIP-strip

We used PIP^TM^ Strip Membranes (Molecular Probes/Invitrogen), with a protocol recommended by the manufacturer. The strip was blocked with 25mL of TBS-T +3% fatty acid free-BSA (EMD Millipore Corporation) for one hour and was then incubated in 5mL TBS-T + 3% fatty acid free BSA containing either GST-RRC-1 (1-150), or GST at a concentration of 0.5μg/mL overnight at 4°C. The membrane was then washed 3 times for 10 minutes each time with 20 mL of the TBS-T+3% BSA. We then incubated the strip with anti-GST (mouse monoclonal 26H1 from Cell Signaling Technology) at 1:1,000 dilution in TBS-T+3%BSA for 1 hour at room temperature. This was followed by 3, 10-minute washes, and then incubation with anti-mouse-HRP (GE Healthcare) at 1:5,000 dilution, and then finally 3, 10-minute washes all with TBS-T+3% BSA. The reactions were detected by ECL and exposure to film.

### *C. elegans* strains

*C. elegans* were grown on NGM plates utilizing standard methods and maintained at 20^0^C (Brenner 1974). The following strains were used:

Wild type, Bristol (N2)
PHX4499, *rrc-1(syb4499)*, by CRISPR which expresses RRC-1 with a C-terminal HA tag (described in Moody et al., 2024)
PHX10067*, rrc-1(syb4499 syb10067)*, by CRISPR, described below, which expresses RRC-1 with a C-terminal HA tag and an in-frame deletion of the PX domain (residues 36-143).
GB390, *rrc-1(syb4499)* after outcrossing 4X to wild type
GB391, *rrc-1(syb4499 syb10067)* after outcrossing 4X to wild type
GB377: *sfEx77* [myo-3p::RRC-1::HA; sur-5::GFP], overexpression in wild type background of full-length RRC-1-HA in muscle from an extrachromosomal array. This strain was created by crossing in the extrachromosomal array [myo-3p::RRC-1::HA; sur-5::GFP], from strain GB372 (Moody et al., 2024) which had been used to rescue *rrc-1(ok1747*), into wild type.
GB378: *sfEx79* [myo-3p::ΔNterm-RRC-1::HA; sur-5::GFP], overexpression in wild type background of in-frame deleted PX domain-RRC-1-HA from an extrachromosomal array. Generation of this transgenic strain is described below.
GB379: *sfIs28* [myo-3p::RRC-1::HA; sur-5::GFP], integrated array (from GB377) overexpressing in wild type background full-length RRC-1-HA in muscle. The integration procedure is described below.
GB380: *sfIs29* [myo-3p::ΔNterm-RRC-1::HA; sur-5::GFP], integrated array (from GB378) overexpressing in wild type background PX domain-deleted RRC-1-HA. The integration procedure is described below.
GB399: GB379 (*sfIs28* [myo-3p::RRC-1::HA; sur-5::GFP]), after outcrossing 3X to wild type.
GB400: GB380 (*sfIs29* [myo-3p::ΔNterm-RRC-1::HA; sur-5::GFP]), after outcrossing 3X to wild type.

#### Generation of GB378 (transgenic overexpression of PX-deleted RRC-1)

To create pPD95.86-ΔPX-RRC-1-HA (deleted 1-140 aa), cDNA fragment encoding residues 141-759 was amplified using a full length rrc-1 cDNA (Moody et al., 2024) and the following primers:

rrc-1-10: SmaI-NheI-rrc-1a ATG-141-
GCGCCCGGGGCTAGCATGCTGAAATTTTTAGAAATTGAC
rrc-1-3: --(EcoRI)-HindIII
CGCAAGCTTCTCTGAATATTCGATTGAATTC

The amplified fragment was cloned into the SmaI and HindIII sites of pBluescript KS+ and a clone was identified by Sanger sequencing to be error-free was called pBS-RRC-1-103. EcoRI, HindIII fragment of pBS-RRC-1-49HA (Moody et al, 2024) was cloned into pBS-RRC-1-103 EcoRI, HindIII sites, resulting pBS-DPX-RRC-1-HA. NheI fragment of pBS-DPX-RRC-1-HA was cloned into pPD95.86 NheI site, resulting pPD95.86-ΔPX-RRC-1-HA. To create GB378, pPD95.86-ΔPX-RRC-1-HA together with pTG96 (sur-5::gfp transformation marker) at a ratio of 1:10 was injected into wild type N2 strain, and screened for the presence of sur-5-GFP expression.

#### Integration of transgenic arrays

Extrachromosomal arrays in strains GB377 and GB378 were integrated randomly into the genome by ultraviolet (UV) irradiation following the procedure of (Mitani 1995) with some modifications (Peter Barret and Jenna Hill, personal communication). Briefly, 40 GFP-positive transgenic L4 hermaphrodites were picked onto 8 OP50 freshly seeded 6 cm plates, 5 worms onto each plate. The plates were then mutagenized in a UV crosslinker at a dose of 250 mJ/cm^2^ for 30 seconds of exposure with the lid of the plates off. Plates were then sealed with parafilm and stored at 20^°^C until starvation estimated to be in 2 weeks. Following starvation, the plates were chunked to new plates and numbered to track each integrant. A total of 11 chunked plates were made for each integrant (full-length RRC-1 and ΔNterminus_RRC-1). These plates were than stored in 20^°^C and allowed to grow for 2 days. From these plates a total of 100, L4 or earlier GFP+ transgenic worms were picked onto separate plates. After allowing to grow for 1 generation, plates were identified that had 100% of the progeny being GFP+, and this transmission rate continued through the F2 generation. One such plate was identified for each extrachromosomal array.

#### Generation of PHX10067

The CRISPR/Cas9 procedures were caried out by SunyBiotech (www.sunybiotech.com). Details about the sgRNAs and repair template used are given in **Supplementary Figure 1**.

### Worm locomotion assays

Swimming and crawling assays were performed on day 2 adults as described previously (Moody et al., 2024).

### RNAi knockdown of *git-1*

RNAi knockdown of *git-1* in GB390 and GB391 was performed as described previously (Moody et al., 2024).

### Western blots

The method of Hannak et al. (2002) was used to prepare total protein lysates from the CRISPR lines, *rrc-1(syb4499)* [RRC-1-HA], [ΔPX RRC-HA], and the transgenic lines that overexpress in a wild type background, RRC-1-HA and ΔPX RRC-1-HA. Equal amounts of total protein were separated on 10% polyacrylamide-SDS-Laemmli gels, transferred to nitrocellulose membranes, reacted with anti-HA (rabbit monoclonal) at 1:1000 dilution and then reacted with goat anti-rabbit immunoglobulin G conjugated to HRP (GE Healthcare) at 1:10,000 dilution, and visualized by ECL reagents.

#### Fixation, Immunostaining, and Confocal Microscopy of Body Wall Muscle

Adult worms were fixed and immunostained using previously described methods (Nonet et al. 1993; Wilson et al., 2012). Primary antibodies were anti-HA (rabbit monoclonal C29F4 from Cell Signaling Technology) used at a 1:200 dilution, and the following antibodies used at 1:100 dilution: anti-PAT-6 (rat polyclonal; Warner et al. 2013); anti-UNC-95 (rabbit polyclonal Benian-13; Qadota et al., 2007), anti-MHC A (mouse monoclonal 5-6; Miller et al., 1983; purchased from University of Iowa Hybridoma Bank), anti-UNC-89 (rabbit polyclonal EU30; Benian et al., 1996), and anti-ATN-1 (mouse monoclonal MH35; Francis and Waterston, 1991). Secondary antibodies were used at a 1:200 dilution including anti-Rabbit Alexa 488 and anti-Rat Alexa 594 purchased from Invitrogen. Images were captured at room temperature with a Zeiss confocal system (LSM510) equipped with an Axiovert 100M microscope and an Apochromat x63/1.4 numerical aperture oil immersion objective. The color balances were adjusted by using Adobe Photoshop (Adobe, San Jose, CA).

## Results

PX-domains share little sequence similarity, so that the detection of such domains in proteins from sequence data alone can be difficult. Accordingly, to query the possible existence of a PX domain in RRC-1, a sequence-based search of the RCSB Protein Data Bank (https://www.rcsb.org) was performed as well as three different protein domain prediction programs: ProSite (https://prosite.expasy.org), PFAM (http://pfam.xfam.org), and NCBI Conserved Domains (https://www.ncbi.nlm.nih.gov/Structure/cdd/wrpsb.cgi). Of these approaches, only NCBI Conserved Domains predicted a PX domain in RRC-1, located at residues 41-146 (Figure 1A). To validate that the identified sequence segment was compatible with a PX domain, we calculated its predicted 3D-fold using AlphaFold3 (https://alphafoldserver.com; Abramson et al., 2024). The predicted model exhibited a high plDDT score (> 90) in all regions other than loops, which indicates high prediction confidence (Supplementary Figure 2). The model displayed a fold consisting of three antiparallel β-strands (β1–β3) followed by three α-helices (α1–α3) (Figure 1B), as is typical of canonical PX-domains (Chandra and Collins 2018). This showed that the identified sequence segment in RRC-1 is compatible with the structurally conserved 3D-fold of PX domains. Next, to determine whether the RRC-1 PX domain possesses functional features characteristic of PX domains, we assessed the presence of the canonical PtdIns3P-binding site, the polyproline motif ΦPxxPxK (Φ: hydrophobic residue), and the secondary PIP2/PIP3-binding site, as described for PX domains (Chandra et al., 2019). The canonical PtdIns3P-binding site was largely absent, with a single amino acid - R38 - remaining conserved (Figure 1B, yellow; Supplementary Figure 3) and no polyproline sequence motif could be detected. However, the secondary PIP2/PIP3 binding site, which typically consists of a surface patch of basic amino acids vicinal to the primary PtdIns3P site, can be predictably matched to the presence of residues R48, H51, R52, R57, R58, H59, and R61 in that region of the fold (Figure 1B, pink: Supplementary Figure 3). This hints at the possibility that PX-RRC1 might have broad specificity for phospholipids and binds a range of these ligands.

**Figure 1.**
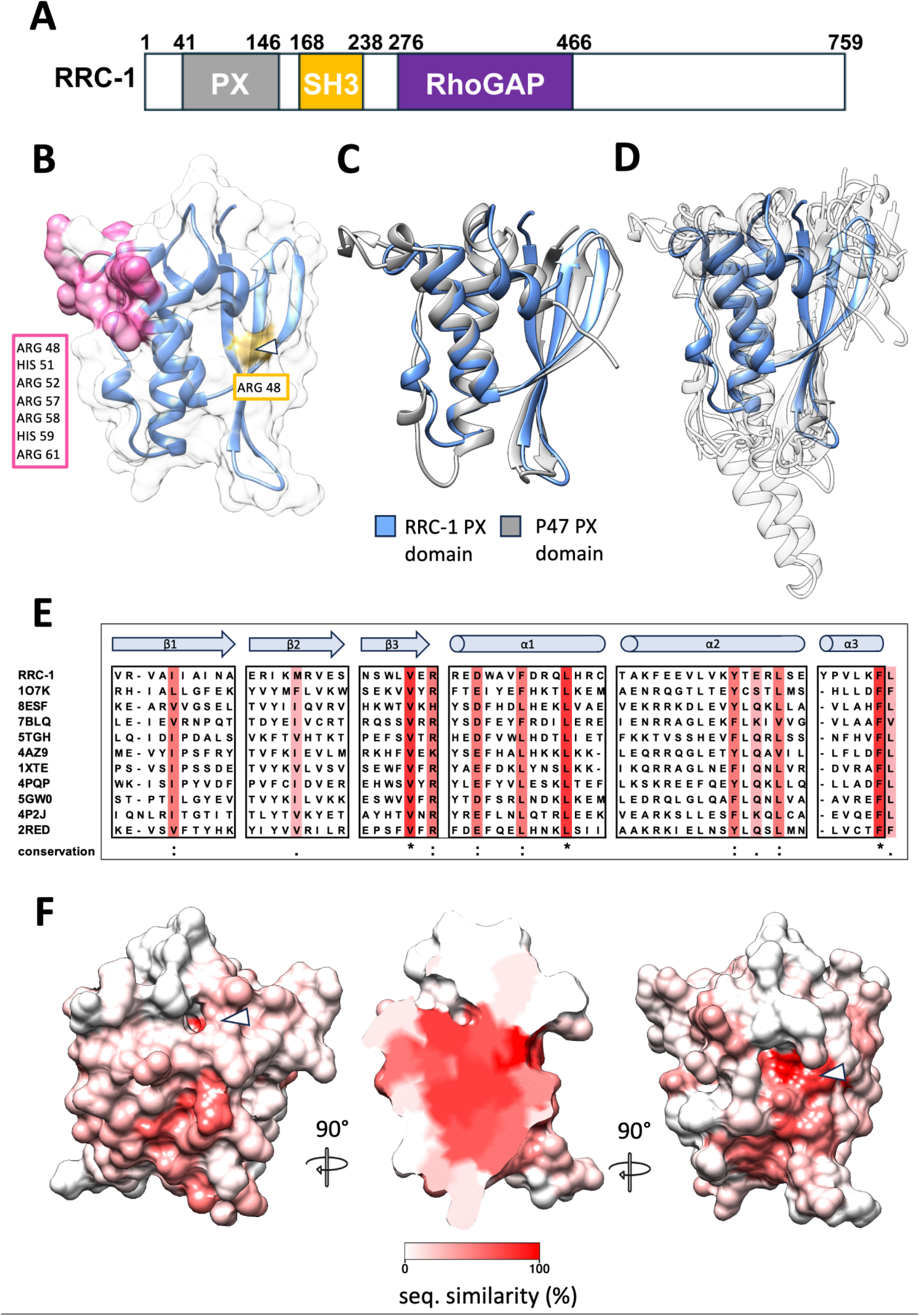
RRC-1 is predicted to comprise a canonical PX domain. (A). Schematic representation of domains in RRC-1 predicted by NCBI Conserved Domains; (B) Predicted 3D-model of the RRC-1 PX-domain displayed both as ribbon representation and molecular surface. The surface representation is colored to show the cluster of basic amino acids at the secondary PIP2/PIP3 binding site (pink) and residue R38, part of the canonical PtdIns3P-binding site (yellow); (C) The 3D-model of PX-RRC-1(blue) is superimposed onto the experimental structure of the PX domain from p47phox (grey), the closest structural homologue. (D) Superimposition of PX-RRC-1 (blue) and the top ten closest PX domains identified using DALI (Holm, 2022) (transparent ribbons). Despite low levels of sequence similarity, the main fold topology is strongly conserved with main differences being only present in loop regions. (E) Structure-based sequence alignment of the secondary structure elements of the PX folds. For each sequence, PDB codes are provided. Conserved residues are coloured in a white-to-red gradient according to increasing conservation levels. (The complete structure-based sequence alignment is provided as Supplementary Figure 2). (F) RRC-1 PX domain colored by conservation as in E. Conserved residues cluster on a deep cavity of the fold (indicated with a white arrow head).

To further confirm its PX-fold identity, we aimed to identify additional sequence determinants of the PX fold in the domain from RRC-1. For this, and since sequence-based approaches did not prove successful in identifying homology, we compared the calculated 3D-model to experimental structures of PX-domains available at the PDB using DALI as search tool (http://ekhidna2.biocenter.helsinki.fi/dali; Holm, 2022) and utilized the identified structural homologues to reveal conserved sequence loci. The closest structural homologue of the RRC-1 PX domain identified in this way was the PX domain from p47phox that displayed only minor structural differences (RMSD=1.8 Å over 91 atom pairs) (Figure 1C). To identify sequence conservation, the top ten hits of the DALI search (Z-score > 8; RMSD < 3 Å) corresponding to the homologues most closely related to PX-RRC-1 were selected for comparison (see Supplementary Table 1). The 3D-model of PX-RRC-1 and the 10 experimental structures so selected were superimposed, revealing a conserved fold with deviations only in loop regions (Figure 1D). A structure-based sequence alignment of the superimposed structures that used UCSF Chimera (https://www.cgl.ucsf.edu/chimera; Pettersen et al. 2004) revealed low sequence identity levels (11–21%) for any given pair (Supplementary Figure 3; Supplementary Table 1). Yet, several hydrophobic residues could be observed to be highly conserved (Figure 1E). The conserved residues so identified showed a strong agreement with conserved residues previously reported to be characteristic of most PX domains (Seet and Hong, 2006), bringing support to the PX identity of the RRC-1 domain. Subsequently, we mapped this conservation onto the 3D-model of the PX-RRC-1 domain and found that the conserved amino acids correspond largely to the protein hydrophobic core as well as forming a small cluster on the protein surface, localizing at a topographical deep cavity that penetrates the fold (Figure 1F). Notably, residue R38 (Figure 1B), part of the canonical PtdIns3P binding site, is located in this conserved surface patch, at the entry to the cavity. Whether this locus still acts as a phosphoinositide-binding site despite lacking complete PtdIns3P binding features is yet to be revealed. Deep cavities comparable to those modelled in PX-RRC-1 are observed in experimental 3D-structures of other PX-domains, e.g. in sorting nexin 3 (PDB entry 7BLQ) and sorting nexin 14 (PDB 4PQP). So far, no function has been assigned to such cavities and their potential contribution to phosphoinositide binding is unknown. Taken together, our bioinformatics analyses lead us to conclude that RRC-1 contains a PX domain of broad phosphoinositide binding specificity.

To experimentally confirm the ability of the RRC-1 PX domain to bind to phosphoinositides, we used a blot (“PIP strip”) containing spots of various phosphoinositides to determine if the PX domain of RRC-1 can bind to phosphoinositides, and if so, which ones. We purified a bacterially expressed GST fusion protein that contains residues 1-150 of RRC-1 (**Figure 2a**) to test for lipid binding. **Figure 2b** displays the results of incubating the strip with GST-RRC-1 (1-150), followed by detection with anti-GST antibodies. There is some binding to the following: phosphatidylinositol 3,5-diphosphate, phosphatidylinositol 3-phosphate, phosphatidylinositol 4,5-diphosphate, phosphatidyl 4-phosphate, phosphatidylinositol 3,4,5-triphosphate, phosphatidylinositol 5-phosphate, and phosphatidyl-serine. This was compared to a control PIP strip which was incubated with GST protein with no resultant binding to any of the phosphoinositides present (**Figure 2b**). We conclude that the predicted PX domain of RRC-1 does have affinity for membrane-localized phosphoinositides.

**Figure 2.**
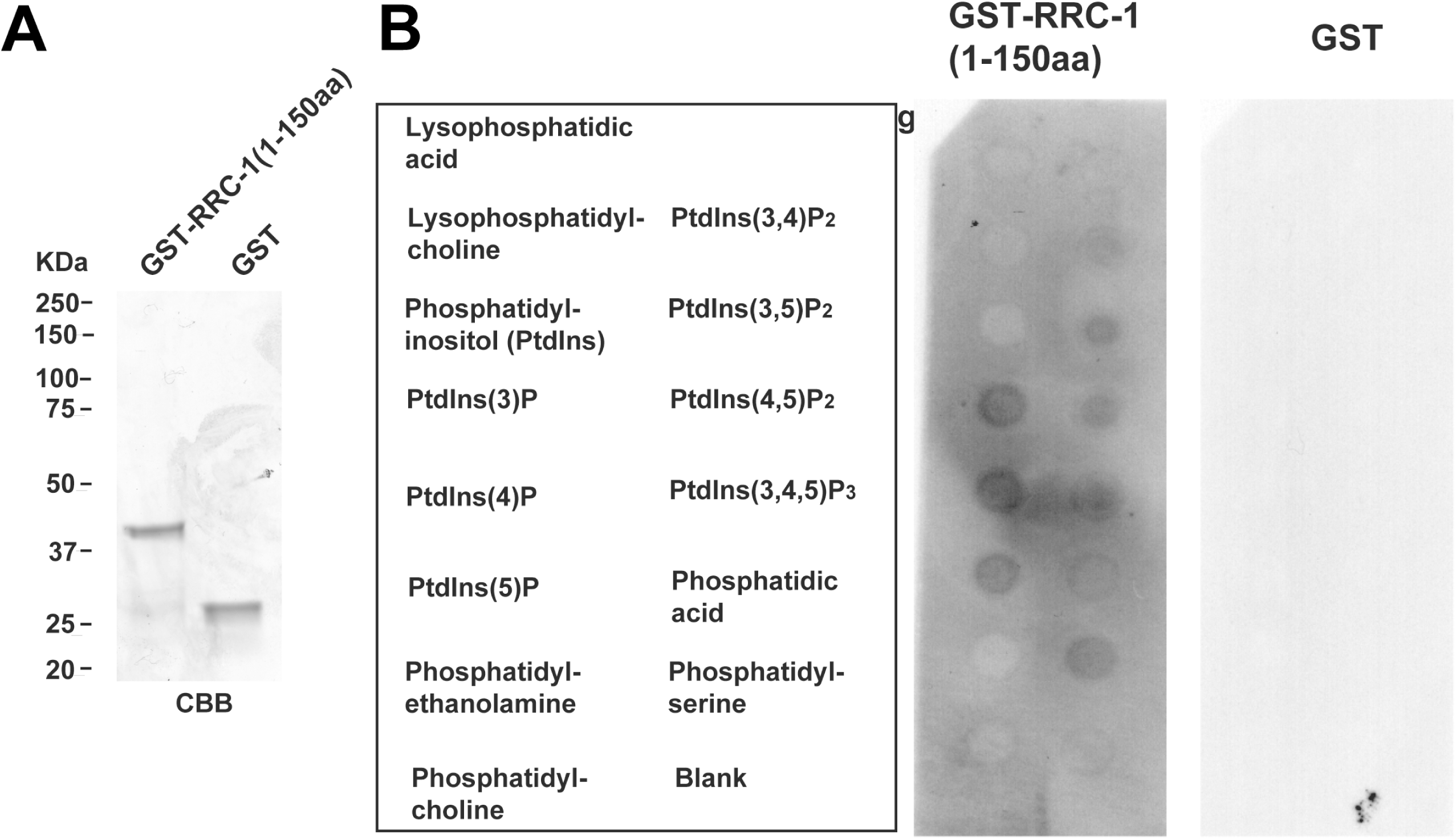
Protein-lipid overlay assay demonstrates that RRC-1 (1-150) binds to phosphoinositides. (A). Coomassie stained SDS-PAGE showing 2 μg of bacterially expressed GST-RRC-1 (1-150), and GST itself. Positions of molecular weight size markers are indicated. (B) Results of incubating GST-RRC-1(1-150) and GST with a PIP Thin Strip (ThermoFisher), and detection using antibodies to GST. Identities of spotted lipids are shown to the left.

To determine the function of RRC-1’s PX domain, we used CRISPR to delete in-frame the coding sequence for the PX domain. This was done in our previously reported (Moody et al. 2024) CRISPR strain that expresses RRC-1 with an HA tag at its C-terminus, *rrc-1(syb4499)*. As shown in **Figure 3a**, we observe expression of ΔPX RRC-1-HA, and it is of expected size. Also, the level of ΔPX RRC-1-HA is only slightly reduced compared to the level of the full length RRC-1-HA (**Figure 3B**). Worms expressing ΔPX RRC-1-HA show the same locomotion as worms expressing full length RRC-1-HA (Figure 4C), which itself did not show a different motility than wild type (Moody et al. 2024). This contrasts with intragenic deletions of *rrc-1* that display reduced swimming and crawling speeds (Moody et al. 2024). Similarly, in contrast to intragenic deletions of *rrc-1* which result in sarcomere disorganization and lack of accumulation of IAC components at the MCB (Moody et al. 2024), ΔPX-RRC-1-HA displays normal sarcomere organization and IAC localization at the MCB (Figure 4D and E).

**Figure 3.**
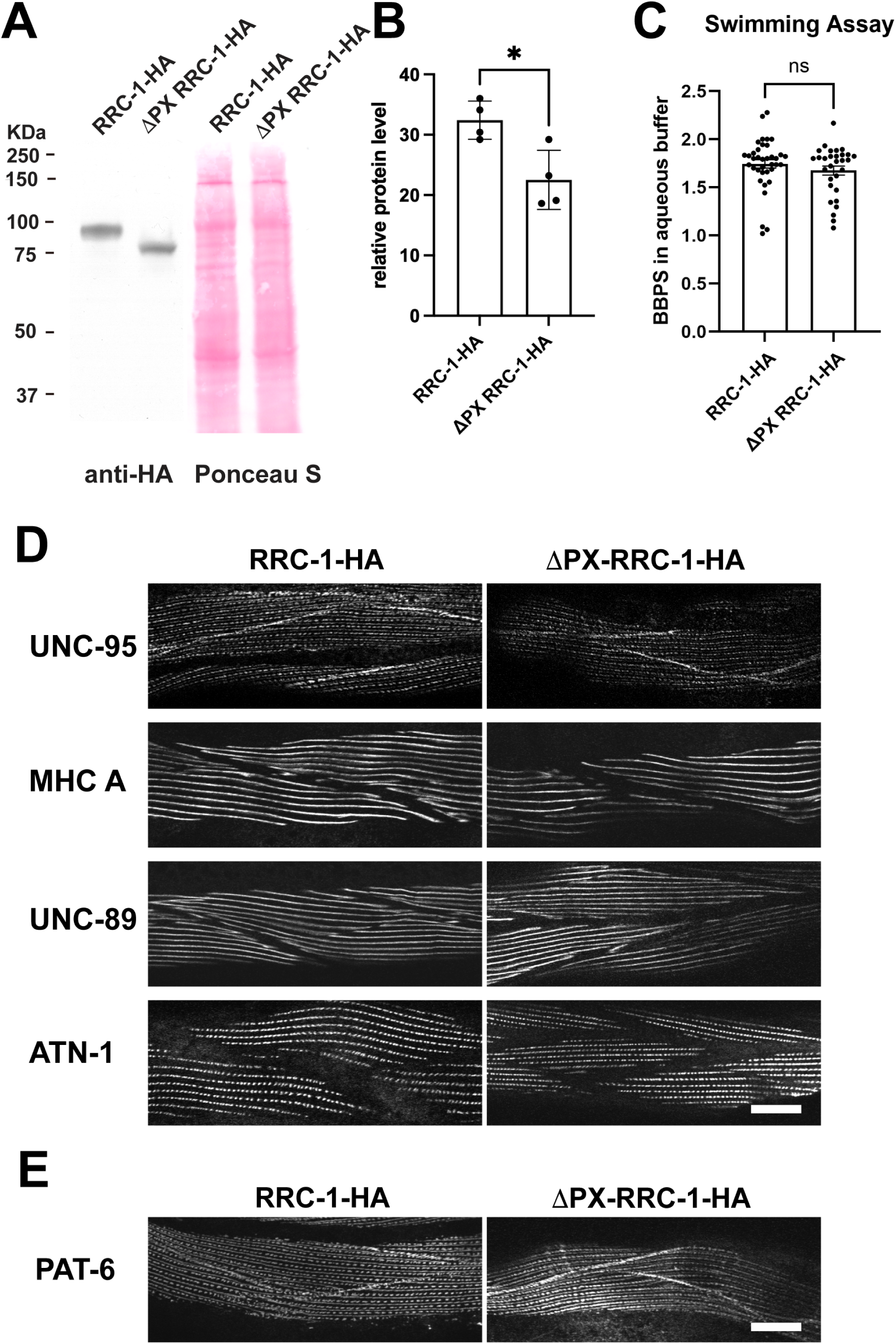
In-frame deletion of the PX domain of RRC-1 results in a normal muscle phenotype. (A). Western blot demonstrating expression of full-length RRC-1 with an HA tag (RRC-1-HA), and RRC-1 with an in-frame deletion of the PX domain (ΔPX RRC-1-HA), both strains generated by CRISPR/Cas9. (B) By quantitative western blot, ΔPX-RRC-1-HA has a slightly lower level of expression than does full length RRC-1-HA. *: p=0.0187. (C) In a swimming assay, ΔPX RRC-1-HA worms swim as well as RRC-1-HA worms, which swim as well as wild type worms (Moody et al. 2024). (D) Immunostaining with antibodies to various sarcomeric proteins shows that DPX-RRC-1-HA worms have normal sarcomere organization, similar to RRC-1-HA worms. Scale bar, 10 μm. (E) The IAC component PAT-6 (α-parvin) localizes normally to MCBs in both DPX-RRC-1-HA and RRC-1-HA worms. Scale bar, 10 μm.

**Figure 4.**
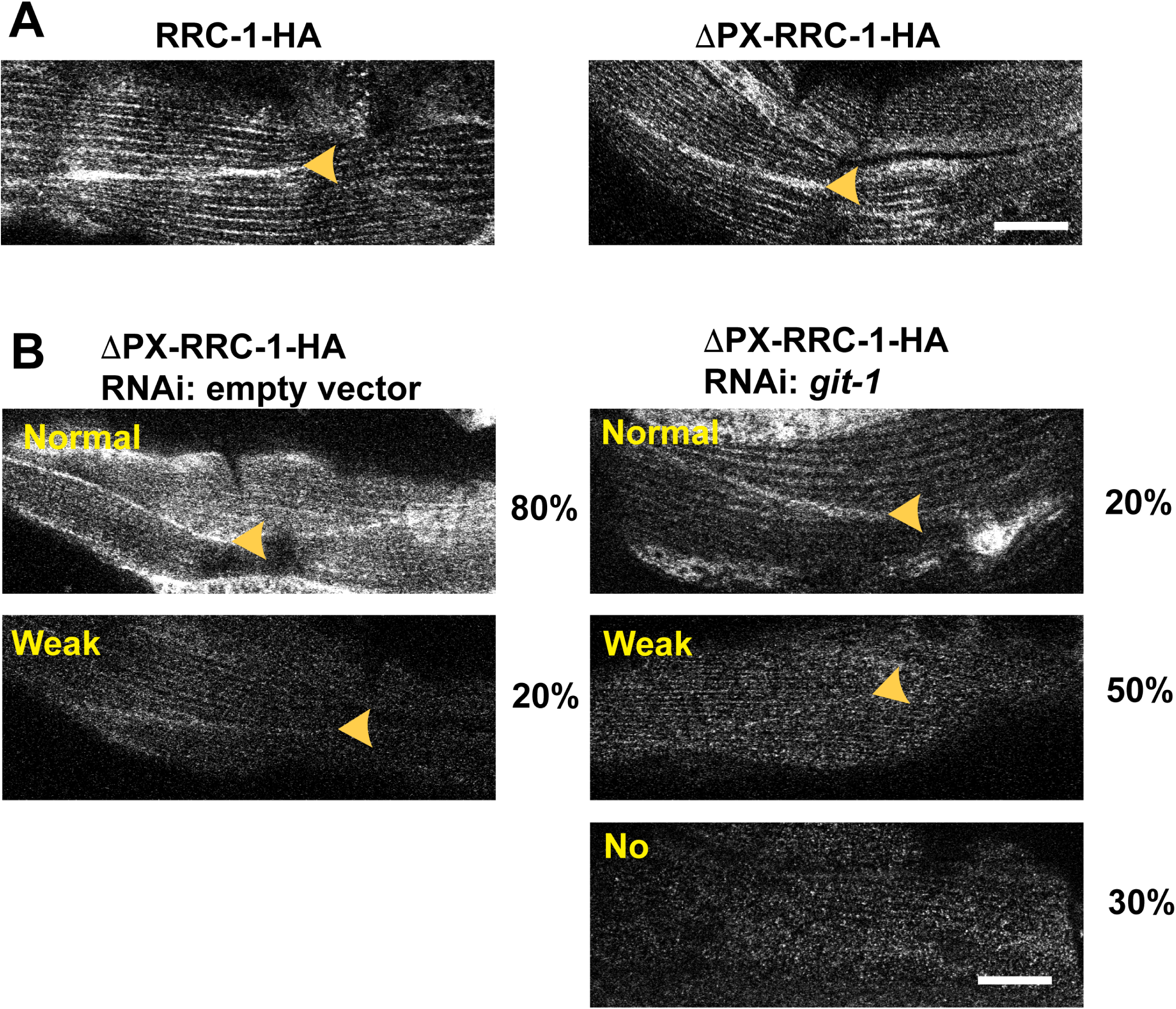
Optimal localization of RRC-1 to the MCB requires both RRC-1’s PX domain and the scaffold protein GIT-1. (A) Nematodes expressing either RRC-1-HA or DPX-RRC-1-HA show normal localization of RRC-1-HA and ΔPX-RRC-1-HA to the MCB, using antibodies to HA. (B) ΔPX-RRC-1-HA subjected to the RNAi empty vector show normal localization of DPX-RRC-1-HA to the MCB at a frequency of 80%, and weak localization at 20% of the animals. However, when ΔPX-RRC-1-HA animals are subjected to *git-1*(RNAi), DPX-RRC-1-HA localizes normally at a frequency of 20%, weakly at a frequency of 50% and not at all at a frequency of 30%. Percentages are based on examining 20 animals. Arrowhead point to MCBs. Scale bars, 10 μm.

ΔPX RRC-1-HA localizes to MCBs and “fuzzily” to dense bodies and M-lines like full length RRC-1-HA (**Figure 4A**). We have reported (Moody et al. 2024) that RRC-1 is part of the PIX complex, which includes the scaffold protein GIT-1. Furthermore, we showed that by RNAi knockdown of *git-1*, GIT-1 is not required for the localization of RRC-1 but is required for the stability of RRC-1. Because we found that the PX domain is not strictly required for RRC-1’s localization to the MCB, and neither is GIT-1 required for RRC-1’s localization to the MCB, we wondered if the PX domain and GIT-1 were redundant for MCB localization. To test this idea, we conduced RNAi for *git-1* in the strain that expresses ΔPX-RRC-1-HA (**Figure 4B**). ΔPX-RRC-1-HA with the RNAi empty vector control showed normal localization to the MCB at a frequency of 80%, and weak localization at 20%. In contrast, ΔPX-RRC-1-HA with *git-1(RNAi)* showed normal localization to the MCB at only 20% frequency, weak localization at 50% and no localization at 30%. Therefore, consistent localization of RRC-1 to the MCB requires both its PX domain and GIT-1.

We have reported that both deficiency and overexpression of PIX-1 results in lack of accumulation of IAC components to the MCB and reduced whole animal locomotion (Moody et al. 2020). We next asked whether overexpression of RRC-1 would have a similar effect and if it depends on the presence of its PX domain. To address this question, we created integrated arrays that overexpress in a wild type background full length RRC-1-HA or ΔPX RRC-1-HA from the nearly body wall muscle-specific *myo-3* promoter. As shown in **Figure 5A**, a western blot shows appropriately-sized proteins are produced from the respective transgenic lines. In addition, the levels of expression are comparable (**Figure 5B**). However, by a swimming locomotion assay, overexpression of full-length but not ΔPX-RRC-1 results in reduced swimming (**Figure 5C**). Consistent with this result, overexpression of full length RRC-1-HA results in reduced accumulation of two IAC components, PAT-6 (α-parvin) and UNC-95 (**Figure 5D**). However, overexpression of ΔPX RRC-1-HA shows normal accumulation of PAT-6 and UNC-95 to the MCB (**Figure 5D**). These results suggest that indeed the PX domain of RRC-1 has a function in muscle, and that likely the membrane localization of RRC-1 via its PX domain is critical.

**Figure 5.**
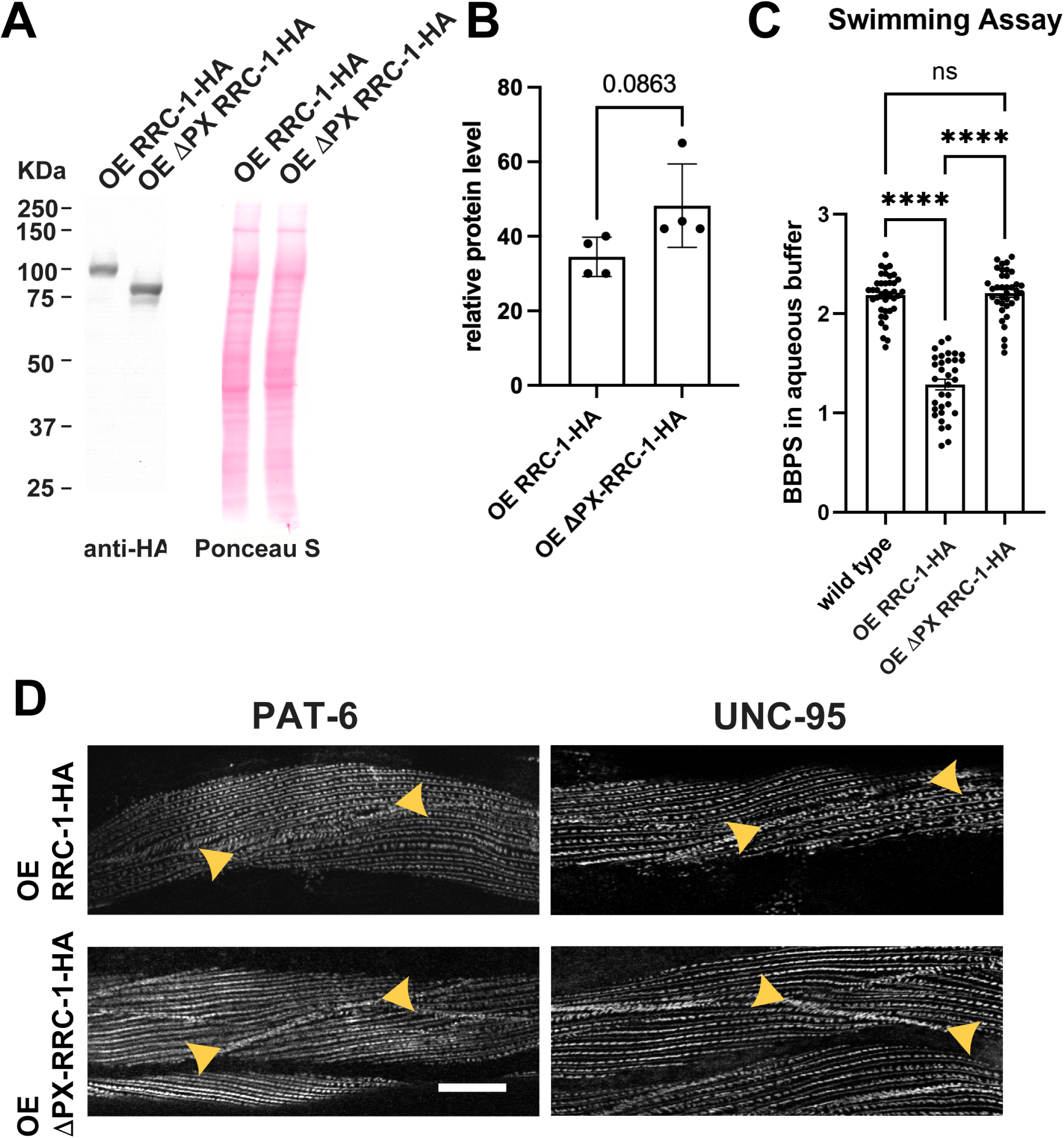
Overexpression of full-length RRC-1 but not RRC-1 with an in-frame deletion of its PX domain results in reduced accumulation of IAC components at the MCB and reduced whole animal locomotion. (A) Western blot demonstrating transgenic overexpression of full-length RRC-1 (OE RRC-1-HA) or RRC-1 with an in-frame deletion of its PX domain (OE ΔPX RRC-1-HA) driven from the nearly muscle-specific promoter from the *myo-3* gene. (B) Western blots reacted with anti-HA indicate no significant difference in the levels of expression from these two transgenes (p=0.0863). (C) Swimming assays show that overexpression of full-length RRC-1 but not RRC-1 with internal deletion of its PX domain results in reduced whole animal locomotion. The indicated transgenes were integrated into the genome by UV irradiation and outcrossed 3X to wild type prior to conducting the swimming assays. Statistical significance was assessed by Dunnett’s T3 multiple comparisons test. ****: p<0.0001; ns: not significant. (D) Overexpression of full-length RRC-1 but not RRC-1 with internal deletion of its PX domain results in decreased accumulation at the MCB of two IAC proteins, PAT-6 and UNC-95. Shown are images of several body wall muscle cells immunostained with antibodies to either PAT-6 or UNC-95. Arrowheads denote portions of MCBs. Scale bar, 10 μm.

## Discussion

We have reported that RRC-1 is required for IAC assembly, and that the RRC-1 protein is likely a component of at least one type of IAC because it localizes to muscle cell boundaries (MCBs) and exists in a complex with another IAC complex protein, PIX-1 (Moody et al. 2024). In the current study, our prediction of a PX domain in RRC-1 (**Figure 1**), and our in vitro binding experiment (**Figure 2**) support the idea that this PX domain binds to phosphoinositides (PIPs). In addition to the PIP strip assay, we sought an additional way to demonstrate PIP binding using a liposome pelleting assay with 6His-PX and liposomes containing as much as 40% PIP3 (3,4,5). We could detect binding, but just above background. We tested the idea that the minimal binding was due to the PX domain already having lipids bound to it, having picked them up during expression and purification from E. coli. Mass spectrometry confirmed our speculation: 6His-PX was found to be associated with several lipids including phosphatidylethanolamine (34:1, and 36:2), and phosphatidylglycerol (32:1, and 34:1)(data not shown). At the least, these results demonstrate that the PX domain of RRC-1 binds to phospholipids.

Our results are further evidence for the importance of PIPs and their metabolism in IAC assembly. Several IAC components in mammals, all of which have nematode orthologs, bind to PIPs, and this is required for their membrane localization and assembly into IACs. For example, kindlin binds to PIP2 and PIP3 through its PH domain, and at least for kindlin 3, with higher affinity to PIP3 (Liu et al. 2011; Ni et al. 2017). Talin binds to PIP2 via its FERM domain (Chinthalapudi et al. 2018). PIP2 is thought to recruit kindlin and talin to the cell membrane and promote localized phase separation of newly forming IACs (Hsu et al. 2023). Vinculin binds to PIP2 via basic amino acids at its C-terminal region (Thompson et al. 2017). The known final component of the PIX pathway, PAK, via a positively charged region binds to PIP2 of the cell membrane and enhances the kinase activity of already activated GTP-Rac or GTP-Cdc42-bound PAK (Strochlic et al. 2010). Recently, we have shown that IPMK-1 (inositol phosphate multikinase), which converts PIP2 to PIP3, is required for normal IAC assembly in *C. elegans* muscle (Sagadiev et al., in revision).

Our results also indicate that the PX domain of RRC-1, although not essential, is still important for RRC-1 function. Analysis of a worm expressing RRC-1 with an in-frame deletion of PX, shows a level of RRC-1 protein that is not appreciably reduced, and displays normal locomotion, normal sarcomere organization and normal localization of IAC components (**Figure 3**). However, RRC-1’s localization to the muscle cell boundary, and likely membrane localization, requires two factors—RRC-1’s PX domain and the PIX complex scaffold protein, GIT-1 (**Figure 4**). We have reported that RRC-1 is a member of the PIX complex of proteins, although we have not demonstrated that RRC-1 binds directly to GIT-1 (Moody et al. 2024). In a wild type background, ΔPX-RRC-1-HA localizes to the MCB and probably the bases of M-lines and dense bodies, just like full length RRC-1-HA (**Figure 4**). Although with low penetrance (∼20%) we observed reduced accumulation of ΔPX-RRC-1-HA in wild type animals, this defect was enhanced to 80% in *git-1* RNAi knockdown animals. This dual requirement for proper RRC-1 localization likely is via membrane association—via the PX domain of RRC-1 and via the ARFGAP domain of GIT-1, since ARFGAP domains may bind to PIP3 (Czech, 2000). The fact that overexpression of full-length RRC-1 but not RRC-1 with an in-frame deletion of its PX domain results in reduced accumulation of IAC components at the MCB and reduced whole animal locomotion (**Figure 5**), further underscores the functional importance of RRC-1’s PX domain. There are two possibilities for the phenotype resulting from overexpression of the full-length RRC-1: (1) A titration or mass action effect--too much of the PIP-binding PX domain could compete out the binding to PIPs and membrane localization of key IAC components like kindlin, talin and vinculin. (2) Too much RacGAP activity: With an intact PX domain, the extra RRC-1 localizes to the muscle cell membrane and there is consequently increased RacGAP activity locally, which results in reduced levels of active GTP bound Rac, similar to loss of function of the RacGEF PIX-1 (Moody et al. 2020).

## Data availability

Worm and bacterial strains, and plasmids are available upon request. The authors affirm that all data necessary for confirming the conclusions of the article are present within the article, figures, and table.

Supplemental material is available at G3 online.

## Acknowledgments

We thank Lizzy Draganova (Emory University) for helpful discussions about lipid binding experiments, and Kristal Maner-Smith at the Emory Integrative Metabolomics and Lipidomics Core for the mass spectrometry of the recombinant PX domain. We thank Andrew Fire (Stanford University) for the *myo-3* promoter plasmid pPD95.86. Wild type *C. elegans*, Bristol (N2), was obtained from the Caenorhabditis Genetics Center, which is funded by the National Institutes of Health Office of Research Infrastructure Programs (P40OD010440). Molecular graphics and analyses were performed with UCSF Chimera, developed by the Resource for Biocomputing, Visualization, and Informatics at the University of California, San Francisco, with support from NIH P41-GM103311.

## Supplementary Material

**Supplementary Figure 1. Generation of strain PHX10067 [rrc-1(syb4499 syb10067)] which expresses RRC-1 with a C-terminal HA tag and has an in-frame deletion of the coding sequence for the PX domain.** This strain was created by SunyBiotech, beginning with PHX4499 which expresses full-length RRC-1 with a C-terminal HA tag (Moody et al. 2020).

**Supplementary Figure 2. AlphaFold model of the RRC-1 PX domain.** 3D-fold of the RRC-1 PX domain (residue 41-146) predicted by Alphafold3. The top model (model B) is shown as a coloured ribbon, where colour corresponds to the per-residue pLDDT confidence score. The subsequent best 5 models are depicted as transparent ribbons. A pLDDT score above 90 (dark blue) indicates high confidence in the predicted model.

**Supplementary Figure 3. Structure-based multiple sequence alignment of the RRc-1 PX domain and its ten closest structural homologues reveals conserved residues in low sequence similarity structures.** The proteins used for the structural alignment are listed with their respective PDB code. Residue boundaries for each sequence are given in brackets. Nomenclature: CIS = cytokine-independent survival; PI = phosphatidylinositol; H = Homo sapiens; M = Mus musculus; T = Thermochaetoides thermophila). Secondary structure elements are marked. Conservation score is given as: * = all residues in that column are identical; : = conserved substitutions;. = semi-conserved substitution. Residues potentially forming the secondary PIP2/PIP3-binding site are highlighted in pink, while the single arginine corresponding to the canonical PtdIns3P-binding site is highlighted in yellow, as described by Chandra et al. (2019)

**Supplementary Table 1. Top ten structurally similar PDB entries to the predicted 3D-model of PX-RRC-1 domain as identified by DALI server.** The first column provides protein names (CIS = cytokine-independent survival; PI = phosphatidylinositol). The second column indicates the species of origin for each protein (H = Homo sapiens, M = Mus musculus, and T = Thermochaetoides thermophile). The third column lists the corresponding PDB codes of the crystal structures. The fourth column presents the DALI Z-scores, indicating the quality of the structural matches. The fifth column shows the mean root-mean-square deviation (RMSD) of the structural alignments. The sixth column reports the length of the aligned region, and the seventh column gives the total sequence length of each protein. The final column indicates the sequence identity (%) from pairwise alignments as output by DALI.

